# Phylogenetic relationship between birds and their magneto-microbiome

**DOI:** 10.64898/2026.01.20.700517

**Authors:** Moshe Leibovitch, Robert Fitak, Eviatar Natan, Yoni Vortman

## Abstract

Animals from a wide taxonomic range can sense earth’s magnetic field, however the underlying mechanism remains one of sensory-biology greatest mysteries. One hypothesis suggests that Magnetotactic bacteria (MTB) serve as the underlying mechanism. This hypothesis predicts that MTB will be detected in animal microbiomes and might show a phylogenetic relationship with their hosts. We examined the phylogenetic relationship between various MTB species across 4,048 avian species using databases of MTB genetic presence across the tree of life and an avian phylogenetic tree. We documented 12 genera of MTB in association with 185 avian species. Three genera, *Magnetospirillum, Magnetovibrio* and *Solidesulfovibrio*, were found at relative high prevalence of positive samples (84%, 33%, 12% respectively). Further, *Magnetospirillum* showed a significant phylogenetic relationship with avian species in general and specifically within Psittaciformes, and Passeriformes. Our results demonstrate the power of harnessing the newly published MTB-database, with specific host-related queries. This analysis, to the best of our knowledge has never been done, and could be replicated across the animal kingdom. The relationship detected suggests an evolutionary and ecological relationship between MTB and avian hosts. These results are consistent with the symbiotic magnetic sensing hypothesis and highlights the potential role of microbiome in sensory physiology.

## Introduction

The magnetic sense remains one of the mysteries of sensory biology, despite decades of research, the sensor behind the magnetic sense remains an enigma (Johnsen 2017; Nordmann et al. 2017; Lohmann et al. 2022). Different hypotheses have been proposed to describe the underlining mechanism of this sense, namely the radical-pair mechanism (Ritz et al. 2000; Hore and Mouritsen 2016), the magnetite-based mechanism (Kirschvink et al. 2001), symbiotic magnetic sensing (Natan and Vortman 2017; Natan et al. 2020), MagR magnetic protein (Qin et al. 2016) and electromagnetic induction (Nimpf et al. 2019), see(Vortman et al. 2025). None of the above-mentioned hypotheses have been unequivocally proven, and some may be relevant to some organisms while others are not. Several issues have probably contributed to this scientific stasis, among those stands out a methodological challenge, clearly isolating the magnetic sense from other senses and accordingly detecting the sensory pathway. Specifically, the fact that birds serve as a model system, and these are highly complex organisms, probably adds to this sensory-biology challenge. Second, it seems that the field had multiple competing paradigms which added to the scientific challenge (Vortman et al. 2025). Taken together, despite extensive research and multiple hypotheses, the magnetic sense remains a sense without a known sensor (Nordmann et al. 2017; Vortman et al. 2025). Out of the above-mentioned hypotheses, the most recently raised hypothesis is the symbiotic magnetic sensing hypothesis (Natan and Vortman 2017; Natan et al. 2020). This hypothesis basically suggests that symbiotic magnetic bacteria that reside within hosting organisms align along Earth’s magnetic field and are sensed by the host.

Magnetotactic bacteria (MTB) are a diverse, polyphyletic group of prokaryotes characterized by the presence of a magnetosome, an organelle containing crystals of magnetite or greigite (Bazylinski et al. 2013). Magnetosomes allow MTB to align along or react to the ambient geomagnetic field. MTB are environmentally ubiquitous and are gathering increasing attention for their unique ecological traits, role in extreme environments, and potential applications in technology and astrobiology (Goswami et al. 2022; Shen et al. 2023). The possibility that MTB may underly the mechanism behind animal magnetic sensing (Natan and Vortman 2017; Natan et al. 2020), was originally criticized because no evidence existed for the presence of MTB in symbiosis with a host animal (Monteil and Lefevre 2020; Leão and Lefèvre 2023). However, recent evidence has documented both the physical presence of MTB in association with a host (i.e., a marine protist (Monteil et al. 2019) and bivalve (Dufour et al. 2014)) and the genetic presence across numerous metazoan taxa (Natan et al. 2020; Fitak 2024). Despite the presence of MTB in association with hosts, conclusive evidence of a symbiosis with a metazoan species has not yet been presented.

Recently, a large, macrogenetic database of MTB presence across metagenomic and animal samples was published (Fitak 2024). This database demonstrated the ubiquity of MTB in various animal samples throughout the animal kingdom and is a valuable addition to the web-based, open databases scientific revolution that is driving science forward. The VertLife database is another powerful example of these web-based resources for researchers (Jetz et al. 2012). The VertLife database facilitates phylogenetic analyses throughout the vertebrate tree of life, allowing simple and rapid creation of specific subsets of vertebrate phylogenetic trees. Here, using these open databases we examined whether MTB detected in birds show a phylogenetic signal. In other words, we hypothesize that if some MTB form symbiotic relationships with birds, then the presence of specific MTB genera will be distributed non-randomly across the avian phylogeny. We further demonstrate the ubiquity of MTB in avian samples and which MTB genera are most abundant and could be targeted for a potential symbiotic relationship with their hosts. We target the class Aves because studies of birds have been essential for navigation research and specifically were the first taxa to have experimental evidence for magnetic sensing (Wiltschko and Wiltschko 1972). Moreover, as flight is in the phylogenetic roots of this group the ability for navigation and possibly to sense the magnetic field is expected to be widespread, currently known in both migratory (Wiltschko and Wiltschko 1972) and resident (Mora et al. 2004) species. Our study further demonstrates how the combination of these two web-based, open databases facilitates novel querying possibilities to examine such phylogenetic relationships throughout vertebrates. The mystery behind the magnetic sense is of a wide taxonomic range, spanning from worms and insects, to reptiles and mammals (Nordmann et al. 2017; Lohmann et al. 2022). Accordingly, we expect future studies to emerge examining the phylogenetic relationship between animals and MTB across various taxa.

## Methods

### Creating the avian magneto-microbiome dataset

The database of the MTB distribution includes the number of DNA sequencing reads from more than 26 million published datasets in the Sequence Read Archive (SRA) that can be unambiguously assigned to 214 MTB taxa (Fitak 2024). We queried this database for the specific datasets derived from avian species (∼150,000 datasets for class = Aves). We observed that the database was heavily biased for domestic chicken samples (*Gallus gallus*) (36% of datasets). Although chicken are known to have a magnetic compass sense (Wiltschko et al. 2007), we omitted datasets originating from *Gallus gallus* due to their marked overrepresentation. We also omitted cultural hybrids, resulting in a cleaned subset of more than ≥90,000 avian samples (hereafter called the “90K” sample database). This 90K sample database was composed of 4,314 differently named avian species. Next, we cross-referenced these species against the VertLife database (Jetz et al. 2012), identifying synonyms, merging subspecies and removing species not existing in the VertLife database. This resulted in 4,048 avian species (supplementary Table 1). For each avian species we averaged the proportion of samples positive for each MTB genus (read count ≥1), creating a final dataset of 4,048 species and 12 MTB genera (supplementary Table 1). We chose to use the MTB genus taxonomic level rather than the species level as there are currently large discrepancies in the assigned species names of many MTB – as they are currently being described in an increasing rate (Natan et al. 2020) – whereas genus is a higher, more stable taxonomic level. Because the dataset was overall quite sparse (many species were assigned 0 counts of most MTB genera) (Fitak 2024), we created a subset of the database that removed all MTB negative samples and included only avian species that had at least one DNA sequencing read assigned across all MTB taxa. This created a subsample of 185 avian species positive for at least one MTB species (see supplementary Table 2).

### Extraction of the avian species’ phylogeny from VertLife

To determine the genetic distances and phylogenetic relationship between avian species we used the avian phylogenetic dataset from the “VertLife” (https://vertlife.org/data/) web phylogenetic database (Jetz et al. 2012). We created 100 phylogenetic trees for both the 4,048 and the 185 species and created consensus trees out of each of the two 100-taxa trees. For each of the two consensus trees we fixed polytomies randomly using functions from the package *ape v5*.*0* (Paradis and Schliep 2019) and *phytools* v2.3 (Revell 2012) package in R. Following the positive relation detected between *Magnetospirillum* and avian species (see results below), we further examined the phylogenetic signal for the presence of *Magnetospirillum* in the three avian orders that had the highest number of positive samples (Psittaciformes, Accipitriformes, Passeriformes). For each of these orders separately we created a phylogenetic consensus tree, using the same protocol.

### Examining the avian phylogenetic signal of MTB detected in avian samples

We took two approaches to examine the phylogenetic relationship between birds and the genera of MTB present in their samples. While the proportion of positive samples is continuous data, the large number of zeros and the difficulty in establishing true negatives (with high probability of missing a positive sample) are better modeled binomially, i.e. whether the species had any positive samples [1] or not [0]. Further, as there is a correlation between number of samples and the probability of detecting a positive sample, we created a binary approach in which species which had 30% of their samples positive were marked as MTB positive and species which had less than 30% of their samples positive were marked as MTB negative, correcting for sample size bias. The 30% threshold was approximately the median proportion of the positive samples (excluding negative samples). We then examined the phylogenetic relationship of the continuous data using Pagel’s lambda (Pagel 1999) and the binary data using the D metric (Fritz and Purvis 2010). Both the Pagel’s lambda and the D metric are indexes of phylogenetic signal, a measure of how closely related species also resemble one another in other traits as expected by Brownian motion along the phylogenetic tree branches as opposed to random distribution along the tree. Pagel’s lambda is an index for continuous data and typically ranges between 0-1, 1 indicating a strong phylogenetic signal (Pagel 1999). While Pagel’s Lamda was developed to analyze continuous data, it is often used also for analyzing discrete data and specifically to incorporate the phylogenetic distances as a covariate in a general model (Pearse et al. 2025). The D metric was created for discrete and rare data where values above 1 indicate an over-dispersed distribution, 1 indicates random distribution, 0 indicates Brownian distribution and negative values indicate an extremely clumped distribution (Fritz and Purvis 2010).

To further strengthen the phylogenetic relationship and confirm that is unlikely due to random chance (or indirect phylogenetic relationship), we examined whether the presence of MTB is associated with habitat type and migratory behavior as covariates – two ecological traits related to flight and geography. Existing trait data were obtained from the Avonet database (Tobias et al. 2022). We performed logistic regression analyses to determine whether these traits predicted the presence of MTB, adding branch lengths as covariates to account for evolutionary relatedness. First, “lambda” (λ) was calculated on the raw bacterial presence data within each order, as mentioned above. Prior to the second calculation, migration habits and habitat data were converted into numerical factors. Subsequently, a linear regression (lm, R version 4.4.1) of bacterial presence against these migration and habitat factors was performed. The residuals of this regression were then used as “corrected” data for the second phylogenetic signal calculation. This approach allowed for the estimation of the phylogenetic signal of bacterial presence while controlling for the influence of ecological factors, thereby isolating the unique contribution of phylogenetic relatedness.

## Results

We examined 150K samples and following basic filtering (see methods), we ended up with a dataset of 90K samples taken from 4,048 different avian species from 44 different taxonomic orders (see supplementary Table S1). Overall, 185 avian species from 20 different orders were positive for at least one of 12 different MTB genera found across all samples (Figure 1). Due to sparsity of the dataset, we examined both the entire (4,048 species) sample and the subset of all MTB-positive species (185 species). The phylogenetic analysis revealed significant phylogenetic signal for three genera of MTB (*Magnetospirillum, Magnetaquicoccus* and *Magnetobacterium*) across the avian phylogenetic tree (Figure 1, Table 1). However, when taking a binomial approach (positive or negative for MTB) and accounting for only cases that 30% of the samples were true positives, only the genus *Magnetospirillum* showed a significant phylogenetic relationship with Aves (Figure 1, Figure 2, Table 1). Other genera showed non-significant phylogenetic signal across this avian phylogenetic tree (or were entirely absent from the sample) when using this approach (counting only >30% positives per species) due to lack of positive samples (Table 1).

**Table 1:**
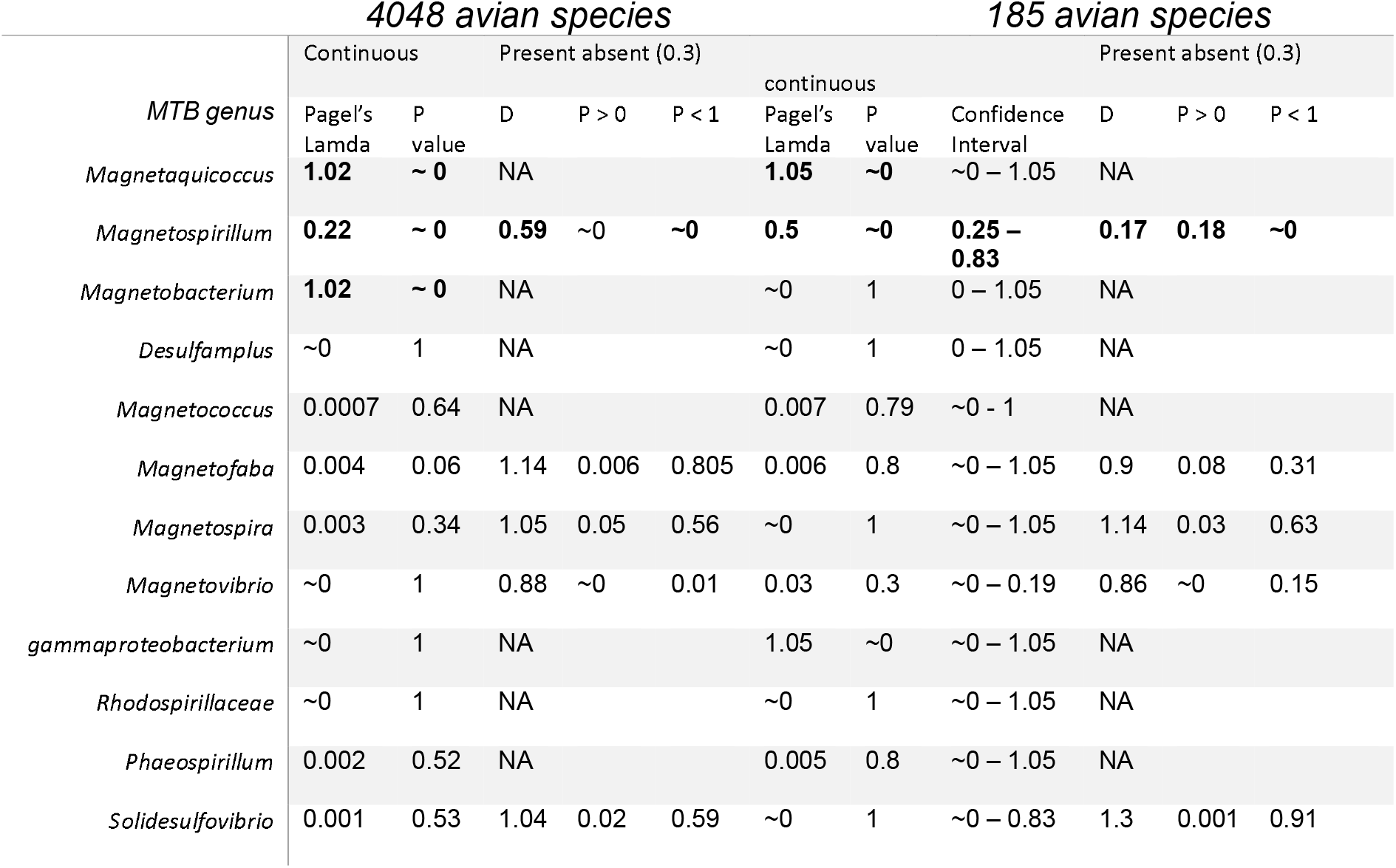
Results for the phylogenetic analysis of each of the MTB genera present in the dataset, and the different analytical approaches. Only *Magnetospirillum*, which is the most abundant in the dataset, shows a significant relationship in all analytical approaches. Computing lamda confidence interval was computationally possible only for the smaller (only positive) subsample. ∼0 denotes P<<0.0001.

**Figure 1.**
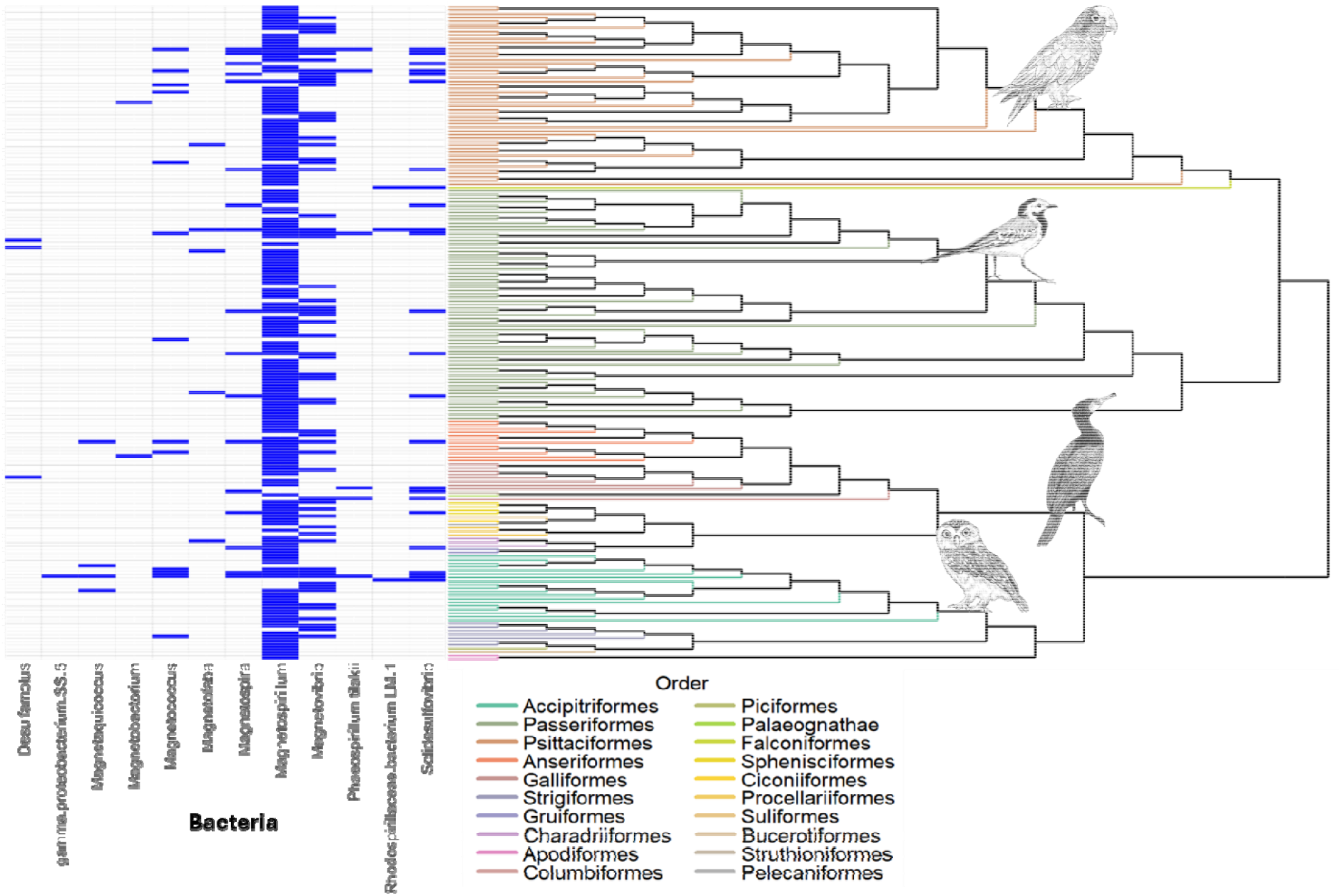
Presence of MTB genera in avian species samples and their phylogenetic relationship. Phylogenetic tree of 185 avian species which were positive for MTB and a Heatmap of the 12 genera of MTB that were detected in the samples (blue = present). This subsample may indicate species that were better sampled or represented in the SRA database, accordingly this subsample was also separately analyzed for the MTB-Aves phylogenetic relationship (Table 1, columns under “185 avian species”). Avian drawing and their taxonomic order from top to bottom: *Pionus senilis* (Psittaciformes), *Motacilla alba* (Passeriformes), *Phalacrocorax carbo* (Suliformes), *Athene brama* (Strigiformes). Drawings by xxx.

**Figure 2.**
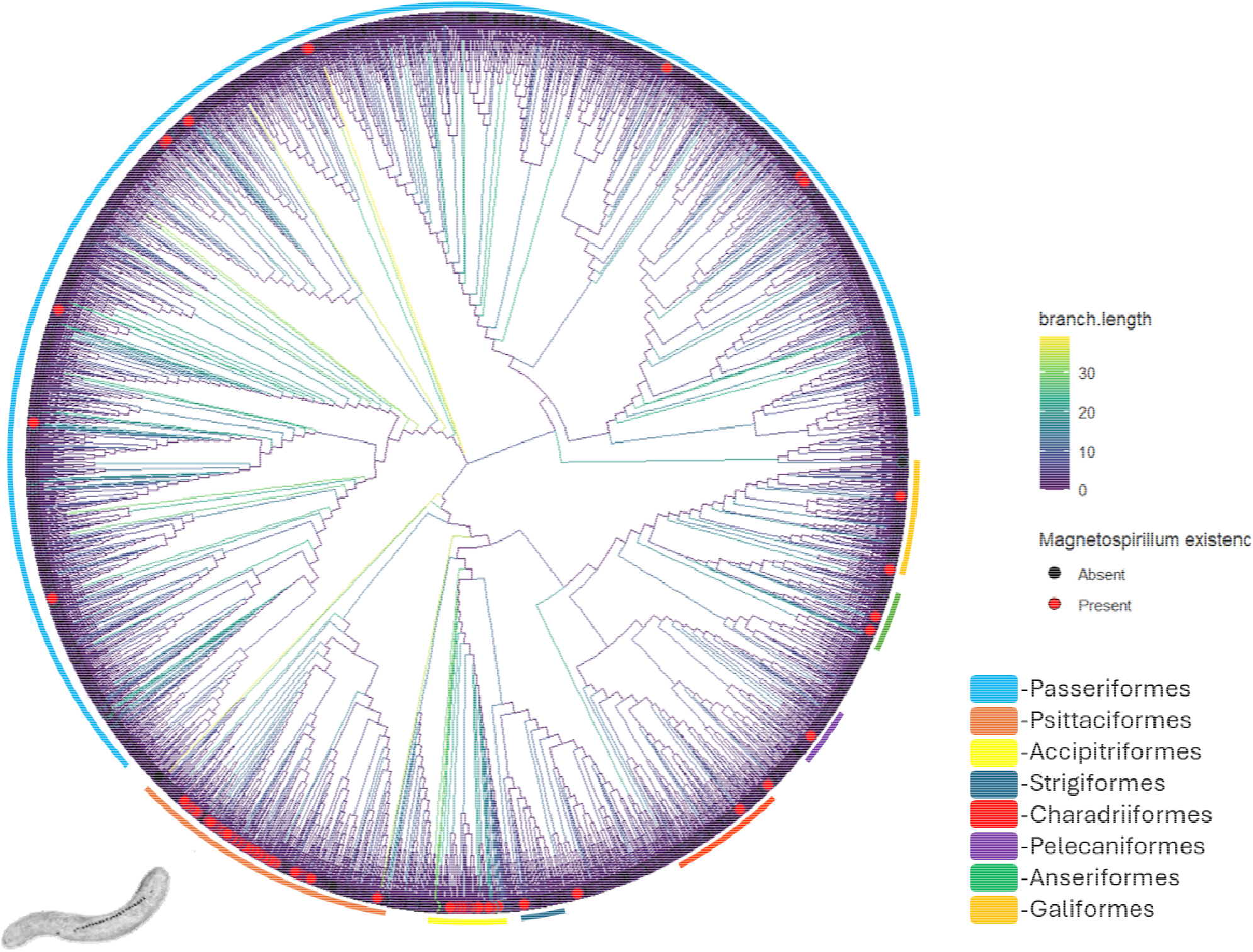
Phylogenetic tree of 4,048 avian species and the presence of *Magnetospirillum* in more than 30% of their sample. *Magnetospirillum*-positive species are indicated by red dots. Branch length is indicated by branch color. It is evident that *Magnetospirillium* is detected more in parrots (*Psittaciformes*) and raptors (Accipitriformes) samples (∼13% and 10% out of these orders’ entire sample respectively), branch length approximately indicates million years ago. Absent indicates not detected in the samples. *Magnetospirillium* artwork: xxx.

Following the detected phylogenetic relationship between *Magnetospirillum* and Aves in general (Figure 2), we further focused on specific orders in which we detected a high number of positive samples, including only species in which at least 30% of samples were positive for *Magnetospirillum*. The three orders in which the sample was high enough to examine any phylogenetic relationship were Psittaciformes, Accipitriformes and Passeriformes. When focusing within each order we detected a significant phylogenetic signal in both Psittaciformes (Figure 3), and Passeriformes but no significant signal in Accipitriformes (Table 2).

**Table 2:** Phylogenetic signal predicting the presence of *Magnetospirillum* within three avian orders. The table shows the estimated phylogenetic signal by itself and incorporated in a model including habitat and migration as co-predicting variables. A species was determined positive for *Magnetospirillum* only if at least 30% of the species samples were positive. Significant values are boldfaced.

| Order | Phylogenetic signal predicting *Magnetospirillum* presence |  | Phylogenetic signal, habitat and migration predicting *Magnetospirillum* presence |  |  |  |  |  |
| --- | --- | --- | --- | --- | --- | --- | --- | --- |
|  | Lamda | P | Lamda | P | Habitat Estimate | P | Migration Estimate | P |
| **Psittaciformes** | **0.40** | **< 0.0001** | **0.39** | **< 0.0001** | -0.006 | 0.603 | 0.089 | 0.248 |
| **Accipitriformes** | ~ 0 | < 0.0001 | ~ 0 | < 0.0001 | **0.029** | **0.004** | -0.006 | 0.857 |
| **Passeriformes** | **0.21** | **< 0.0001** | **0.21** | **< 0.0001** | 0.0009 | 0.089 | 0.002 | 0.204 |

**Figure 3.**
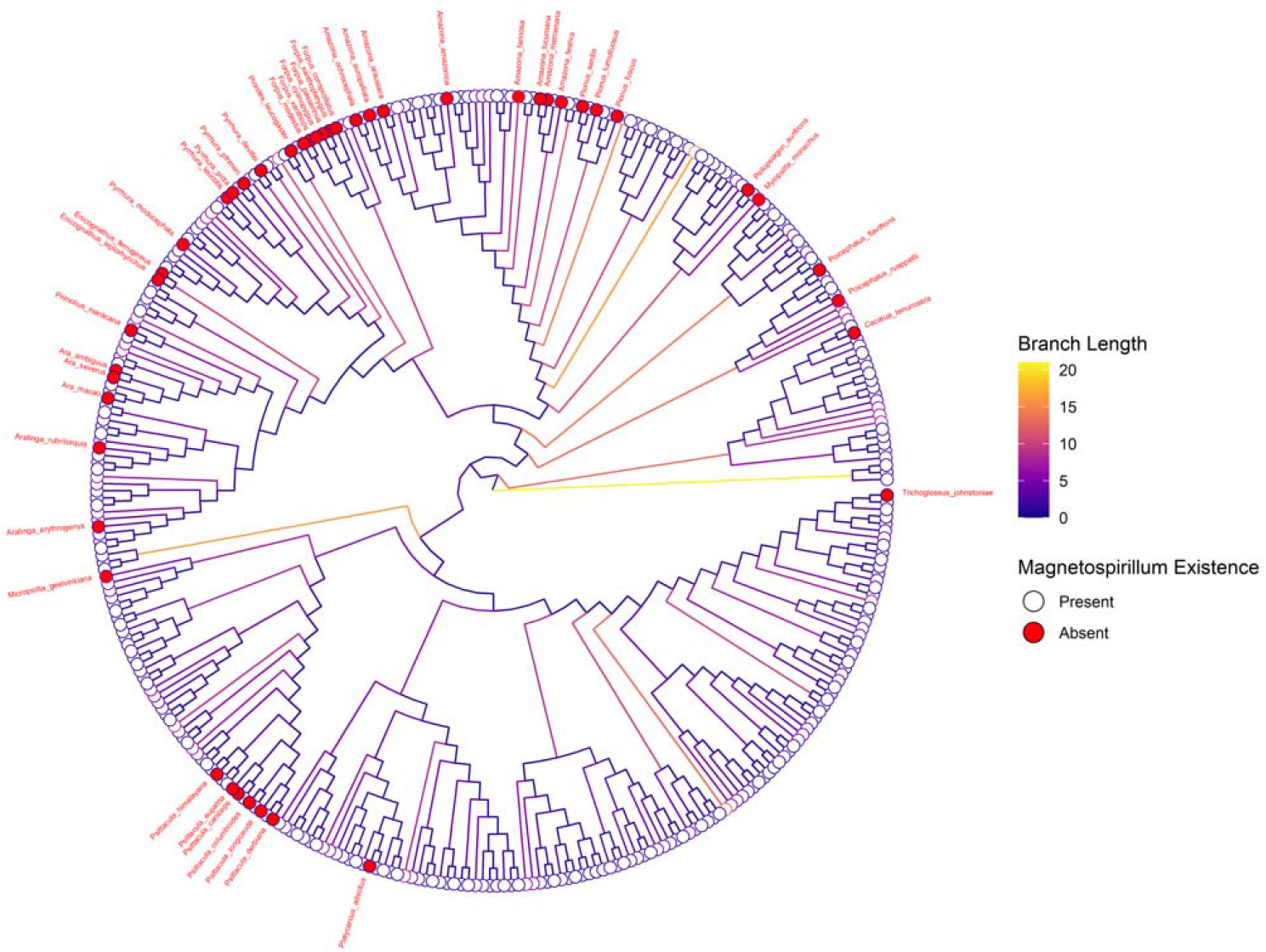
Phylogenetic tree of species from the order Psittaciformes. Species in which at least 30% of their samples were positive for *Magnetospirillum* are indicated with a red dot and the corresponding species name. For detailed statistics see Table 2. Branch lengths are indicated with branch color, approximately indicating million years ago.

## Discussion

Understanding how animals navigate through space is a fundamental aspect of biology (Wynn and Liedvogel 2023). While the ability to sense the magnetic field has been demonstrated in a wide taxonomic range, e.g. insects, fish, amphibians, mammals, reptiles and birds, the magnetic sense remains one of sensory-biology greatest mysteries and keeps drawing broad attention from the scientific community (Nordmann et al. 2017; Johnsen et al. 2020). While several hypotheses regarding the underlying mechanism behind the magnetic sense have been proposed, the symbiotic magnetic sensing hypothesis is in its infancy and accordingly has been relatively less studied (Natan and Vortman 2017; Natan et al. 2020). Using the recently published metagenomic dataset, we report the abundance of MTB within avian samples, a group with relatively less MTB-positive samples in the dataset (Fitak 2024). We further demonstrate that by combining this dataset with existing databases we can examine phylogenetic relationships that may exist between MTB and vertebrate hosts. We found that 12 genera of MTB were present in avian samples and that three genera had a significant phylogenetic signal across bird species. Removing the effect of sample size and taking a more conservative approach still revealed a significant phylogenetic relationship of *Magnetospirillum* with avian species. The phylogenetic relationship between host phylogeny and their MTB microbiome may indicate a long-evolving symbiosis.

Since the symbiotic magnetic sensing hypothesis has been raised, criticism mainly on the probability of detecting MTB in animal samples has been raised (Monteil and Lefevre 2020; Leão and Lefèvre 2023). The current dataset refutes this criticism entirely. The view that detecting MTB in animal samples was unlikely or otherwise incidental contamination may be attributed to the fact that many labs working on MTB were not focusing on animal microbiomes, but rather environmental samples (Lin et al. 2014). The significant phylogenetic signal refutes contamination and supports an MTB–host association. Furthermore, the fact that the first wide support of MTB presence in host came from existing public open databases, demonstrates that this result was waiting out there to be picked up (Natan et al. 2020). Similarly, this publication also exploits the public open magneto-microbiome database which is based on Sequence Read Archive from the National Center for Biotechnology (Fitak 2024), again demonstrating that the presence of MTB in animals’ samples was detected by various researchers and simply never queried. Following the general phylogenetic relationship detected between birds and *Magnetospirillum*, we examined whether this relationship is also apparent within the three orders with the highest number of positive samples using the conservative approach (counting only species that 30% of their samples were positive, as a positive species, Psittaciformes, Accipitriformes, Passeriformes). Both within the Psittaciformes (see Figure 3) and the Passeriformes (Table 2) the presence of *Magnetospirillum* shows a significant phylogenetic signal, further supporting the general trend. Moreover, as a phylogenetic signal between two species could be a result not of direct symbiosis but rather following a geographic relationship which has a common phylogenetic history, we examined within these subsamples whether habitat, migration or phylogeny predicts the distribution of *Magnetospirillum* within these two orders. Even within the combined model nor migration, nor habitat significantly predicted the presence of *Magnetospirillum*, in the orders that demonstrated a significant phylogenetic relationship, in contrast, the phylogenetic signal remains significant (and almost identical) in both analysis methods (Table 2). Unlike Psittaciformes and Passeriformes, in Accipitriformes (which do not show a significant phylogenetic relationship with MTB), habitat significantly predicted presence of *Magnetospirillum*, however the parameter estimate is very low, thus significant but with a low effect. The lack of significant relation between migration estimates and the presence of MTB within the three orders may seem surprising, however, we emphasize two factors. First, as the database is comprised from various sampling efforts done by different researchers and different sampling protocols and subsequent different lab efforts, the data is noisy. Accordingly, we emphasize that negative results should be taken with caution as there is a low signal to noise ratio. Furthermore, the distribution of the samples could be critical in picking such trends, and specifically in the order Psittaciformes the majority of species in our sample are non-migrating species thus it could be an order that is not suited to examine relationships with migration. Second, while migration and navigation are obviously connected, non-migrating species navigate regularly, and accordingly non-migrating species have a magnetic sense (e.g. (Kimchi et al. 2004; Mora et al. 2004; Wiltschko et al. 2007)). The notion that migrating species should have magnetic navigation capabilities while resident species could settle for less is probably biased due to our own navigation capabilities, magnetic sensing is prominent in species which move relatively short distances (e.g. mole rats (Kimchi et al. 2004) and bees (Liang et al. 2016)). Thus, regardless of distance, movement of an organism is a fundamental characteristic of life, it determines the fate of individuals and accordingly orientation mechanisms are expected to evolve also in short distance moving organisms (Nathan et al. 2008). Despite all the abovementioned potential noise in the database, the phylogenetic relationship between Aves and *Magnetospirillum* is detected both in the general sample and within two of the three orders in which we could detect a large enough sample size of positive species to examine, regardless of geography or migration.

It is not surprising that among all MTB genera examined, a significant phylogenetic signal was detected specifically with the genus *Magnetospirillum*. This genus is perhaps the most well-described and with the most known species (Fitak 2024), thus accordingly it is by far the most common MTB genus detected in the avian samples. The lack of significant phylogenetic relationship of the other MTB genera could also be a result of low detection of these genera in the samples, as many fewer species have been described. In addition, negative samples could be due to sampling efforts and amount of microbiome DNA in the sample rather than presence absence of a specific bacteria in the host. The last is especially true for birds as they are very light vertebrates which produce very small amounts of feces, as is portrayed in the low proportion of positive samples of this class with respect to other tetrapods (Fitak 2024) (this is specifically true for Passeriformes). Thus, negative results should be interpreted cautiously, accordingly we think that all genera detected in the avian samples could be considered as potentially important for future studies. Specifically, the genera detected in non-negligible proportions. In addition to *Magnetospirillum*, we further suggest *Magnetaquicoccus Magnetobacterium, Magnetovibrio, Solidesulfovibrio, Magnetospira* and *Magnetococcus* as also potentially important genera. As scientific knowledge of these less-studied taxa grows, detection of these genera in animals and specifically avian samples will likely become easier. Of the various MTB genera examined, one, *Magnetospirillum* was most abundant in general and specifically in parrots (and showed a significant phylogenetic signal within that order).

In our work we examined the presence of specific MTB genera in avian samples and accordingly the phylogenetic relationship of MTB in birds. The significant phylogenetic relationship detected may suggest a “phylosymbiosis”, defined as “Similarities in host-associated microbial community structure that mirror the phylogenetic relationships of the hosts” (Mallott and Amato 2021). Indicators of phylosymbiosis are always intriguing as they suggest that microbiota have shaped host ecology and evolution. As these microbiotas are not present in the host DNA and may be transmitted horizontally or gathered from the environment and not necessarily heritable, such “genetic” relationships suggest a close and reliable dependency across generations. If symbiotic MTB affect the host navigation ability through magnetic sensing, such a scenario could lead to evolving a phylogenetic relationship. Although a mechanism of symbiotic magnetic sensing has yet to be reported experimentally, these indicators of a phylogenetic relationship between birds and MTB seem a promising support for the symbiotic magnetic sensing hypothesis (Natan and Vortman 2017; Natan et al. 2020). However, we emphasize that this result does not demonstrate that these bacteria serve as a magnetic sensor. Animal navigation is a complex multidisciplinary scientific endeavor (Wynn and Liedvogel 2023) because many species have multiple navigation mechanisms and strategies, and for many the magnetic sense is not preferred relative to other sensory cues, making it difficult to experimentally isolate (Ioalè et al. 2006). Specifically, while Aves have been considered a model species for magnetic sensing, the distribution of this sense is not known as they have multiple senses which take part in the navigational task (Mouritsen 2018). The magnetic sense is present in both migratory e.g. (Kishkinev et al. 2015) and resident e.g. (Mora et al. 2004) avian species and also present in a wide taxonomic range (Nordmann et al. 2017). The mystery behind the magnetic sense is relevant across the animal kingdom (see a recent example, Goforth et al. 2025). Specifically the leading hypothesis for bird magnetoreception, the radical-pair hypothesis (Hore and Mouritsen 2016; but see Vortman et al. 2025), could not be relevant for several specific non-avian species such as the blind mole-rat (Kimchi et al. 2004) and turtles that sense the magnetic field in darkness (Lohmann et al. 2001). Thus, it would be intriguing to examine other taxa that are present in the dataset to see whether similar or different MTB genera share a phylogenetic relationship with their host. To the best of our knowledge our reported analysis has never been done and could be easily replicated with other taxa, specifically tetrapods and Chordata classes are also available at VertLife, demonstrating the ease by which these databases can be utilized by researchers. Furthermore, such an analysis could be replicated using other MTB databases or MTB genomic samples. We hope that this work which follows the publication of the magneto-microbiome dataset (Fitak 2024) will inspire similar work in the future. This work does not prove the symbiotic magnetic sensing hypothesis; however, these results are inline with the hypothesis, adding another small piece to the magnetoreception puzzle.

## Data availability statement

For data and supplementary materials please contact authors at:

## Supplemental information

Table S1. Excel file containing 4,048 avian species. For each avian species we averaged the proportion of samples positive for each 12 MTB genera.

Table S2. Excel file containing 185 avian species positive for at least one MTB species Document S1. R code for both phylogenetic analyses

